# Predicting Serotonin Detection with DNA-Carbon Nanotube Sensors Across Multiple Spectral Wavelengths

**DOI:** 10.1101/2024.01.06.574485

**Authors:** Payam Kelich, Jaquesta Adams, Sanghwa Jeong, Nicole Navarro, Markita P. Landry, Lela Vuković

## Abstract

Owing to the value of DNA-wrapped single-walled carbon nanotube (SWNT) based sensors for chemically-specific imaging in biology, we explore machine learning (ML) predictions DNA-SWNT serotonin sensor responsivity as a function of DNA sequence based on the whole SWNT fluorescence spectra. Our analysis reveals the crucial role of DNA sequence in the binding modes of DNA-SWNTs to serotonin, with a smaller influence of SWNT chirality. Regression ML models trained on existing datasets predict the change in the fluorescence emission in response to serotonin, ΔF/F, at over a hundred wavelengths for new DNA-SWNT conjugates, successfully identifying some high- and low-response DNA sequences. Despite successful predictions, we also show that the finite size of the training dataset leads to limitations on prediction accuracy. Nevertheless, incorporating entire spectra into ML models enhances prediction robustness and facilitates the discovery of novel DNA-SWNT sensors. Our approaches show promise for identifying new chemical systems with specific sensing response characteristics, marking a valuable advancement in DNA-based system discovery.

## 1. Introduction

Single-wall carbon nanotubes (SWNTs) represent promising nanomaterials for sensing and imaging a broad variety of biomolecules^1^. Their large potential is attributed to their non-bleaching near-infrared fluorescence emission, which is suitable for analyte detection in a wide range of complex biological samples^2–6^. To be used in sensing applications, SWNTs are often noncovalently functionalized by adsorbed polymers, solubilizing them in aqueous environments through the formation of a “corona phase” on the SWNT surface. Adsorbed polymers create a surface for analyte adsorption, and a diverse range of polymers have been utilized for SWNT functionalization, including nucleic acids, peptides, surfactants, lipids, and peptoids^7,8,17–19,9–16^.

Among the various functionalization approaches, single-stranded DNA-functionalized SWNT conjugates are the most ubiquitous. They have been extensively employed in optical sensing of biologically important small analytes^5,17,20–22^, as well as for polynucleotide delivery in genetic transformation applications^23,24^, and for chirality sorting of multi-chirality SWNT samples into chirality-pure constituents^9,25–29^, or SWNT enantiomer separation^30,31^. In the context of DNA-SWNT conjugates utilized for optically sensing molecular analytes or separating SWNT chiralities, the DNA sequence plays an essential role. The DNA must simultaneously exhibit high affinity binding to both analytes and the underlying SWNT surface. Furthermore, this binding should result in a significant change in the SWNT optical response, ΔF/F, only in the presence of the target analyte. Identifying new DNA-SWNT nanomaterials that exhibit sensitive and specific responses to desired small molecule analytes poses a challenging problem, requiring the development of innovative data science approaches^17^.

Recently, advanced data analytics approaches have been used to predict new nanomaterials with specific biological behaviors^32^. Many of these approaches involve the acquisition and curation of large datasets in experiments, followed by the use of the artificial intelligence (AI) algorithms to understand and predict material properties and functional behaviors based on these datasets. The nanomaterial development and optimization with AI has been done for purposes of developing new nanomedicines and nanomaterials for drug delivery^33,34^, nanomaterials for detection of cancer biomarkers^35,36^, nanomaterials for sensing biologically important analytes^17^ or toxic metabolytes^37^, as well as predicting the interactions of nanomaterials with the complex biological environments, such as nanomaterial biodistribution^38^ or adsorption of proteins to nanomaterial surfaces^39^.

In the context of predicting nanomaterials that serve as optical sensors for molecular analytes, the desired functionality typically involves a discernible change in the emission spectrum as the analyte is introduced. This spectrum provides the intensity of the sensor light emission across various wavelengths, serves as an indicator of the analyte’s concentration. Efficient development of sensing nanomaterials would benefit from the ability to predict complete emission spectra for nanomaterials of diverse compositions. One way to achieve this goal is to experimentally prepare numerous systems with varying compositions, acquire the emission spectra in response to the selected analyte, and use the resulting data to train machine learning (ML) models for predicting new materials with improved emission response to the analyte. So far, ML approaches have demonstrated success in predicting absorption and emission wavelengths and quantum yields for molecules, offering comparable results to traditional methods (e.g. density functional theory calculations) but at a fraction of the computational cost^40^. Neural network-based ML models are also successful at predicting multidimensional optical spectra and fluorescence properties of chromophores in complex environments^41^. Different types of ML models were also adept at predicting the fluorescence emission spectra of nanomaterials used as fluorescent or luminescent probes^42^, as well as for predicting emission spectra of DNA-templated silver nanoclusters, for which the training accuracies were found to be greater than 80%^43^.

In recent years, ML methods have made significant strides in addressing various questions related to DNA-SWNT-based materials. For instance, ML facilitated a systematic exploration of DNA sequences for sorting carbon nanotubes, effectively separating specific chiralities from SWNT samples typically prepared as mixtures of chiralities^44,45^. ML models were also developed using optical signals from DNA-encapsulated quantum-defect-modified SWNTs in serum samples from individuals with ovarian carcinoma and healthy counterparts^36^. These models demonstrated an impressive ability to detect ovarian cancer, achieving 87% sensitivity at 98% specificity when tested on new patient serum samples. Another recent study introduced a DNA-SWCNT–based photoluminescent sensor array, leveraging optical responses to train ML models for detecting gynecologic cancer biomarkers in patient samples and fluids^35^. In our research, we applied ML classification and regression techniques to predict the response of DNA-SWNT sensors to a crucial neurotransmitter, serotonin^17^, whose biological functions in the brain and throughout the human body warrant further investigation. Our ML approaches successfully predicted five new sensors with responses surpassing any of the other DNA-SWNT conjugates in the original dataset.

In our earlier work^17^, ML models were trained solely on the responses of DNA-SWNTs to serotonin at a single wavelength (1195 nm), extracted from a spectrum encompassing optical responses across a range of wavelengths (850 nm – 1340 nm). Notably, the remaining information from the broader spectrum was left untapped in ML predictions. In the current study, we leverage information from multiple wavelengths in the experimental spectra to train models for predicting wavelength-specific ΔF/F responses for DNA-SWNT conjugates featuring new DNA sequences. For each novel sequence, ΔF/F responses are predicted at over a hundred wavelengths. Following the prediction of segments of emission spectra, we then conduct statistical analyses to create a distribution of ΔF/F predictions for a given sequence. This examination allows us to assess the robustness and confidence levels of predictions for sensors with high responses.

## Methods

### 2.1 Dataset preparation and preprocessing

The dataset used to train and test ML models contains the fluorescence emission spectra of DNA-SWNT conjugates with varying DNA sequences before and after the addition of 100 μM serotonin analyte. The spectra were obtained for samples in aqueous solutions, and the experimental conditions were described in detail in Refs.^17,46^. The dataset was assembled from the data for 136 different DNA-SWNT samples, each containing a unique DNA sequence, summarized in **Table S1**. SWNT samples contained SWNTs of different chiralities and DNA molecules had sequences of the type C_6_X_18_C_6_, where X_18_ was the variable part of the sequence. The collected spectra of fluorescence emission reported intensity values at wavelengths in the near infrared range between 850 nm and 1340 nm. For each DNA-SWNT sample, there were spectra of the sample before and after the addition of 100 μM serotonin. The fluorescence response of each DNA-SWNT sample, ΔF/F(λ) (also called simply ΔF/F), was calculated as ΔF/F(λ) = (F(λ)-F_0_(λ))/ F_0_(λ), where F_0_(λ) and F(λ) are the measured fluorescence intensities at a given wavelength λ before and after the addition of serotonin, respectively. Since the experiments were performed in triplicate, the final fluorescence response of each DNA-SWNT sample was chosen to be an average of the triplicate measurements. Out of the total of 136 different DNA-SWNT systems in the original dataset, six systems were reserved for independent validation of our ML models, resulting in a final dataset of 130 DNA-SWNT systems that is used for training and testing the ML models.

Examination of ΔF/F(λ) values for all the DNA-SWNT conjugates in our dataset revealed that some wavelengths are associated with ΔF/F values that are strongly sequence-dependent and span a wide range, whereas other wavelengths are associated with ΔF/F values that are similar for all the examined sequences and span a narrow range (**Figure S1a**). The wavelengths at which ΔF/F values span a narrow range are likely to be unhelpful for training the ML models, and we eliminated them from our dataset using a defined quantitative criterion. This criterion finds the wavelengths (parts of the spectra) for which ΔF/F values span a wide range. For all wavelengths, we determined the maximum and minimum ΔF/F values, 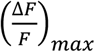 and 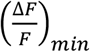, across all 130 DNA-SWNT conjugates present in our dataset. Then, we identified the wavelengths at which the difference between the minimum and maximum ΔF/F values was greater than a defined threshold value t:

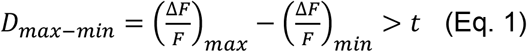

Here, the wavelengths of interest for training the ML models are those for which ΔF/F values span a relatively large range, namely, when D_max-min_ > t = 1.5. With t = 1.5, approximately 30% (312 out of 1025) of the wavelengths are selected for the subsequent training of ML models. **Figure S1a** shows the regions of spectra that are selected for ML model development using the threshold defined in Eq. 1.

Next, we examined the distributions of ΔF/F values at specific wavelengths. Examples of these distributions at several wavelengths are shown in **Figure S2**. We posited that the quality of the ML models is going to be better if the distribution is broader (spanning a larger range of ΔF /F values) and bimodal in nature. The criterion in Eq. 1 already selected the wavelengths that span the larger range of ΔF/F values. Next, we removed some of the DNA-SWNTs with ΔF/F values falling in the middle of the distributions as described below, in order to obtain a more bimodal-like distribution at each wavelength. Increasing the bimodality in distributions leads to better separation in ΔF/F values of high and low response systems: for example, our previous work showed that increasing the bimodality of distributions and increasing the separation in ΔF /F values of two classes of systems (high and low response systems) led to higher f^1^ scores of the classification ML models^17^. The mathematical criterion for removing the sequences from the middle of the distribution was wavelength-specific. At each wavelength *j, λ*_j_, we calculated the median and mean ΔF/F values. The average of the mean and median, *α*, represents the center of the distribution at a given wavelength. The parameters defining the range of ΔF/F values around *α* from which some datapoints will be removed to create a gap in the dataset is depicted in **Figure S1b**. This range has a flexible point around which the sequences will be removed, α’ = α + *imm*, where *imm* is a variable that increases the average of the mean and median and is defined to be one of the values in the set {0, 0,05, 0.1, 0.15, 0.2}. The range also has a flexible width, which is varied by a variable called tolerance, *tol*. The sequences that are removed from the dataset at a given wavelength have ΔF/F value between α’ and the upper threshold, f_upper_, or have ΔF/F between α’ and the lower threshold, f_lower_. The upper and lower thresholds are defined as f_upper_ = α’ + *tol* and f_lower_ = α’ – *tol*. The removal of sequences with ΔF/F between the two thresholds results in a dataset of DNA-SWNTs with high response to serotonin (ΔF/F > f_upper_, shown as a green region in **Figure S1b**) and DNA-SWNTs with low response to serotonin (ΔF/F < f_lower_, shown as a gray region in **Figure S1b**).

### 2.2 Machine Learning Model Training

The preprocessed dataset described above was used to train regression ML models using the Support Vector Machine (SVM) algorithm, which stands out as a favored algorithm in supervised learning, particularly adept at handling scenarios with small sample sizes and high-dimensional data challenges^47^. Each regression model used the DNA sequence as input and predicted ΔF/F at a given wavelength as output. The 18-nucleotide (nt)-long DNA sequences were represented as one-hot encoded (1 x 72) vectors. The DNA sequence vector dimensions were determined by each of the 18 positions in the DNA sequence being occupied by one of four possible nucleotides, A, C, T, and G. The model training procedure was designed to iterate until it identified a predetermined number of regression models. Namely, the procedure continued until it found five models with coefficients of determination, r^2^, surpassing 0.4, when comparing the predicted and the measured ΔF/F values of the testing part of the dataset at a given wavelength.

## 3. Results and Discussion

### 3.1 Fluorescence emission change of DNA-SWNT conjugates in response to serotonin in the 850 nm to 1340 nm wavelength range

Previous experiments collected the fluorescence emission of DNA-SWNT conjugates for 30-nucleotide-long DNA sequences of the type C_6_X_18_C_6_, where X labels variable nucleotides in strands, obtained before and after the addition of the serotonin analyte^17,46^. These samples, and specifically the SWNTs in these samples, are optically active, providing the near infrared fluorescence emission spectra of DNA-SWNT conjugates (**Figure 1a-b**). The collected spectra have multiple peaks of varying intensity, which correspond to emissions by SWNTs of different chiralities present in the experimentally prepared samples. For all 136 DNA-SWNT conjugates in the original dataset, the intensity of the optical emission either stayed the same or increased to a variable extent after the addition of serotonin.

**Figure 1.**
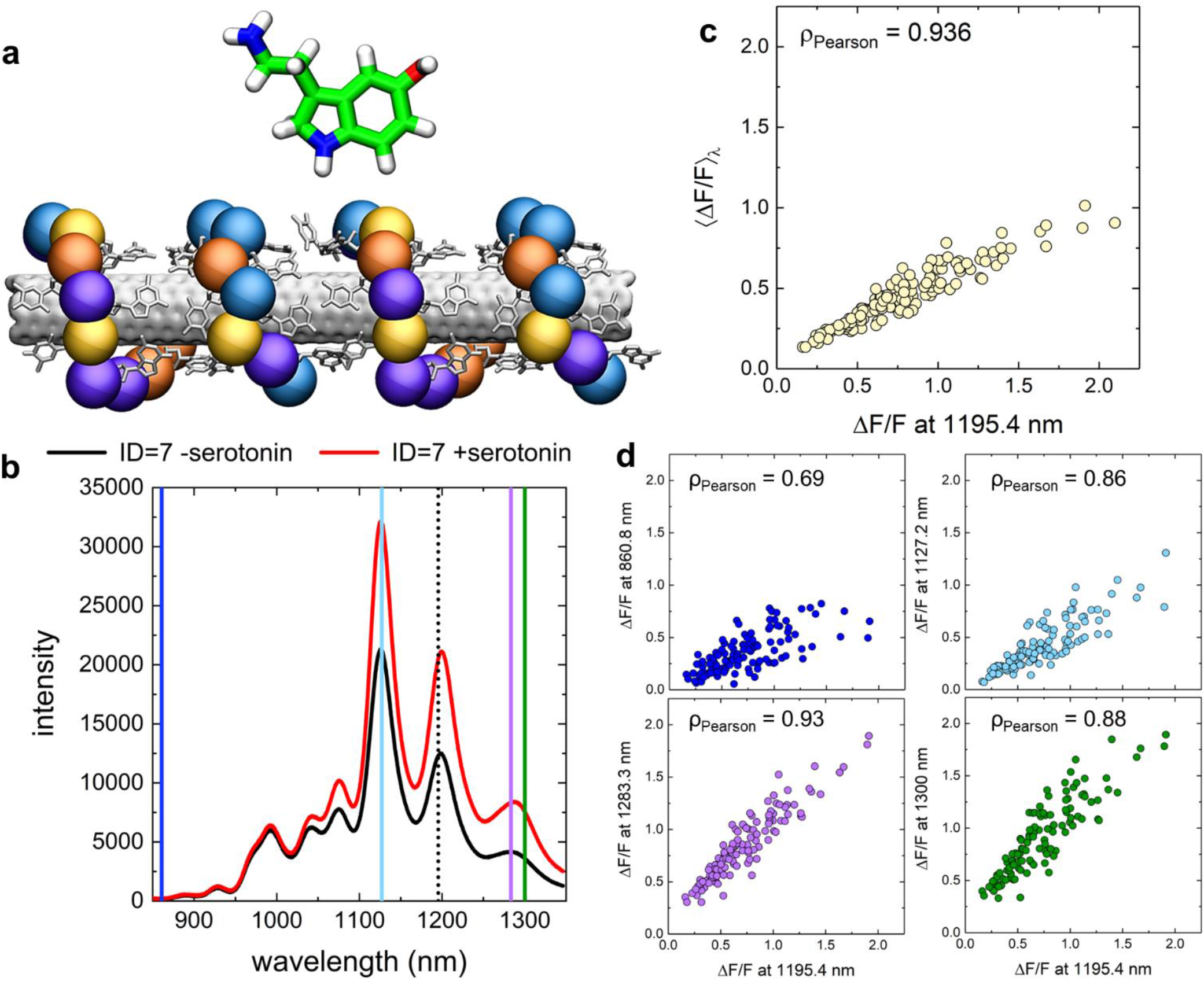
Analysis of relative fluorescence emission changes in response to serotonin analyte for SWNTs of different chiralities in DNA-SWNT conjugates. **a**. The system under investigation. **b**. Example spectra of one of the DNA-SWNT sensors (sequence ID 7). Fluorescence spectra before the addition of serotonin to DNA-SWNT suspension (black trace) and after the addition of 100 μM serotonin (red trace). **c**. Plot of cumulative ΔF/F values and ΔF/F values from 1195 nm wavelength (for (8,6) SWNT) from spectra of selected 96 DNA sequences in response to serotonin. d) Plot of ΔF/F values read at 1195 nm and ΔF/F values at four other wavelengths from spectra of the same 96 DNA sequences in response to serotonin.

In our previous work, our analysis and machine learning efforts were focused on the fluorescence emission at ∼1195 nm center wavelength, which corresponds to the emission of SWNTs with (8,6) chirality. Here, we first examine how the fluorescence emission change (ΔF/F) varies in response to serotonin at specific different wavelengths and for the integrated response over all the wavelengths (cumulative ΔF/F, also labeled as ⟨*ΔF/F*⟩_*λ*_). The plots in **Figure 1c-d** examine the correlations between the fluorescence emission change (ΔF/F) at 1195 nm and either the averaged response over all the wavelengths, ⟨*ΔF/F*⟩_*λ*_, or wavelengths 861 nm, 1127 nm, 1283 nm and 1300 nm. All the plots show strong correlations between ΔF/F at 1195 nm by (8,6) SWNT and the other examined ΔF/F responses. The weakest correlations are observed between ΔF/F values at 1195 nm and 860 nm, with the Pearson coefficient of 0.69, and the strongest correlations are observed between ΔF/F values at 1195 nm and 1283 nm, as well as ⟨*ΔF/F*⟩_*λ*_, where the Pearson coefficients are 0.93 and 0.94, respectively. The observed significant correlations suggest that it is the DNA molecule when adsorbed to the SWNT that primarily determines the analyte binding strength, the extent of the perturbation of the SWNT environment by the analyte, and the intensity of emission by DNA-SWNT samples present in the system. The observation also suggests a possibility that some DNA sequences have a special binding mode to serotonin analyte in the presence of a hydrophobic SWNT surface, since the SWNT chirality plays a less significant role in analyte binding at the SWNT surface and the intensity of DNA-SWNT fluorescence emission.

### 3.2 Predicting fluorescence emission change (ΔF/F) in response to serotonin by DNA-SWNT conjugates at selected wavelengths with ML regression models

Next, ΔF/F response to serotonin of DNA-SWNT conjugates for all the wavelengths across the spectra from previous experimental measurements^17,46^ was used to train machine learning regression models. The workflow of the approach taken in the present work is shown in **Figure 2**. The input dataset for training our models initially contained 136 distinct DNA sequences, which in experimental systems wrap the SWNTs. The 18-nucleotide (nt)-long DNA sequences were represented as one-hot encoded (1 x 72) vectors, and for each DNA sequence, there is an associated matrix of ΔF/F values at all measured wavelengths in the range from 850 nm to 1346 nm.

**Figure 2.**
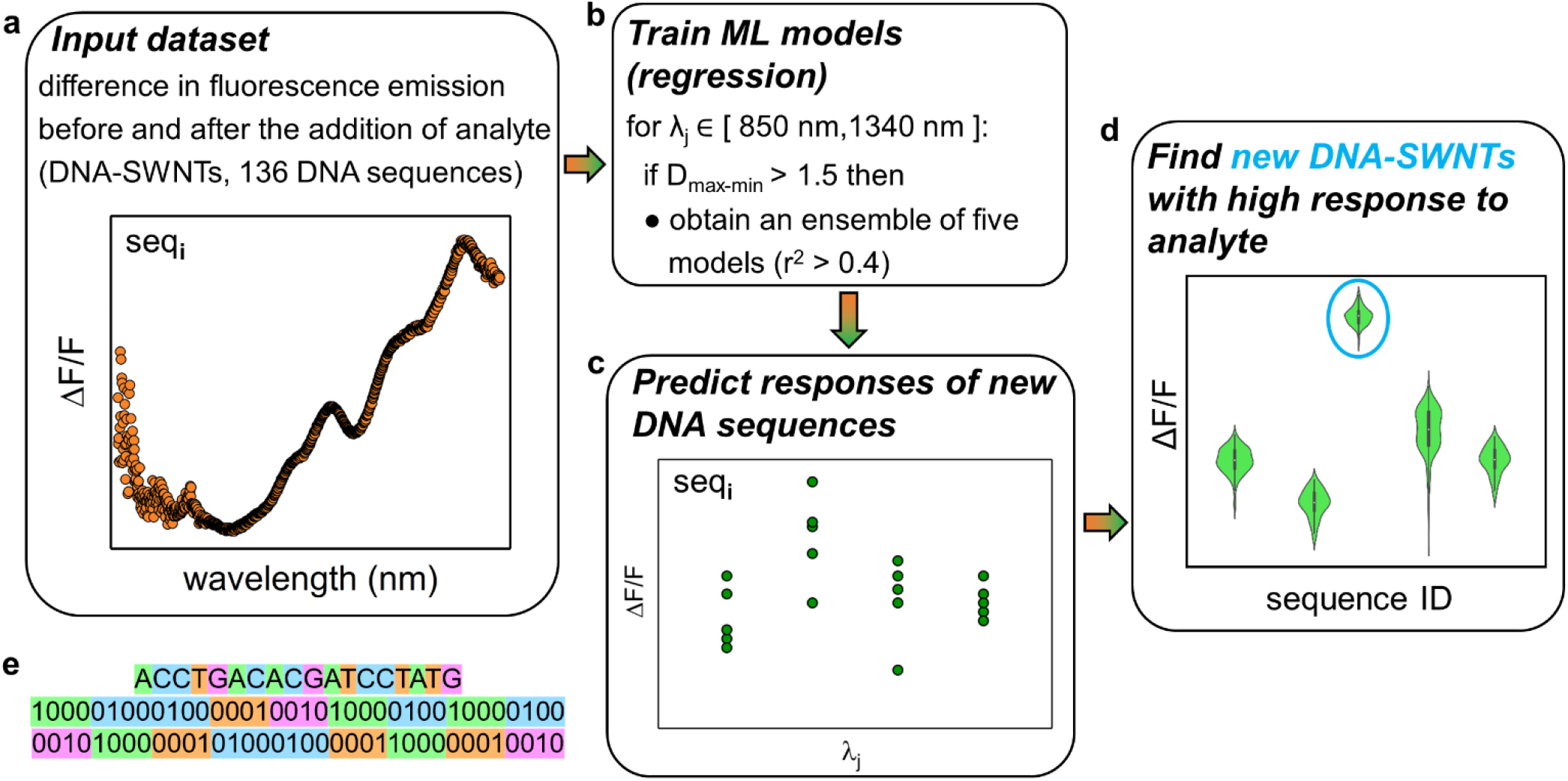
Machine learning workflow for predicting the fluorescence emission change (ΔF/F) of DNA-SWNT conjugates in response to serotonin. a) The input data contains ΔF/F values for wavelengths between 850 nm and 1340 nm for each of 130 DNA-SWNT conjugates experimentally examined in Refs.^17,46^ ΔF/F values were determined from the fluorescence emission spectra before and after the addition of 100 μM of serotonin. b) Using the input dataset, machine learning regression models are trained for each wavelength that has a wide distribution of ΔF/F values, as defined by a quantitative criterion described in Methods. The training is performed on random selections of the 80% of the dataset until at least five models are found for which the coefficient of determination, r^2^, is greater than 0.4 in the analysis of the measured and predicted ΔF/F values in the remaining 20% of the dataset. c) The saved regression models are used to predict ΔF/F values at selected wavelengths for DNA-SWNT conjugates with new DNA sequences. d) The predicted ΔF/F values for DNA-SWNTs with new DNA sequences are statistically analyzed to determine sequences that lead to outlier responses. e) One-hot encoded representation of DNA sequence (ID = 1) from Table S1.

In the next step, we analyzed the distributions of ΔF/F values of all 136 DNA-SWNT conjugates at all wavelengths, one at a time. Example distributions, shown for three wavelengths in **Figure S2**, demonstrate that at some wavelengths, such as λ = 865.3 nm, there are very few ΔF/F values greater than 1, resulting in narrow distributions. However, for other wavelengths, such as λ = 1195.4 nm and λ = 1300.5 nm, the distributions are much broader and have a larger number of ΔF/F values greater than 1 and extending up to the value of 2. We hypothesized that the width of the distribution of ΔF/F values associated with the given wavelength may affect the quality of the machine learning models trained to predict ΔF/F values for new DNA sequences in DNA-SWNT conjugates: the larger variability of ΔF/F responses is assumed to lead to machine learning models that will be better at distinguishing higher response from lower response sequences and thus be more successful. Therefore, we introduced a quantitative criterion to identify wavelengths with wider distributions of ΔF/F values, as described in Methods. The wavelengths for which the criterion was satisfied were then selected as wavelengths for which we train ML models to predict ΔF/F values for new DNA sequences in DNA-SWNT conjugates. Prior to training the models associated with individual wavelengths, some sequences with intermediate values of ΔF/F were removed from the input data, resulting in the input dataset with more bimodal ΔF/F distribution at a given wavelength. Furthermore, six DNA sequences, labeled S1 – S6, were removed from the training dataset of 136 sequences, to perform independent testing of trained models (**Table S2**). Three of these sequences had consistent low ΔF/F response to serotonin and three remaining sequences had consistent high ΔF/F response to serotonin in experiments. After the input data were processed as described, by removing the input data for S1 – S6 sequences, the wavelengths with narrow ΔF/F distributions, and some of the individual datapoints (sequences and their ΔF/F values) from the input data associated with the remaining wavelengths with the goal of achieving bimodal-like distributions, we trained ML regression models to predict ΔF/F values for new DNA sequences in DNA-SWNT conjugates at each retained wavelength. Each model is designed to predict ΔF/F value of a given sequence for a defined wavelength.

For each defined wavelength, we trained many distinct support vector machine regression models (up to 2^32^), until we obtained at least five models for which the experimental ΔF/F and the predicted ΔF/F values resulted in a coefficient of determination r^2^ greater than 0.4 or until the maximum number of models was reached. Our procedure results in five predicted ΔF/F values for each selected wavelength for a new DNA sequence in DNA-SWNT conjugate. Finally, we prepare distribution plots of all the predicted ΔF/F values in the form of violin plots for all the new DNA sequences of interest. These plots are then examined for outlier sequences (with either exceptionally high or exceptionally low distributions of ΔF/F values), which can be suggested for experimental testing.

The predicted fluorescence emission change (ΔF/F) in response to serotonin for selected range of wavelengths for two representative sequences, S1 and S4, are shown in **Figure 3a**. The plotted responses are obtained using one regression SVM model per wavelength. The predicted responses differ from the experimentally measured responses: the predicted ΔF/F values of the low response sequence S1 are overestimates and the predicted ΔF/F values of the high response sequence S4 are underestimates. Furthermore, the differences between the measured and the predicted ΔF/F values are significantly more pronounced for the high response sequence. However, there are also shared trends in the measured and predicted ΔF/F responses, namely, the peaks and valleys in the measured and predicted curves occur at similar wavelengths. We further examined the differences between the measured and the predicted ΔF/F values from ensembles of regression models at all the selected wavelengths (289 wavelengths in total) for all the six test sequences (S1-S6). As shown in **Figure 3b**, the predicted ΔF/F values of low response sequences (S1-S3) are always overestimates by a difference of 0.2 to 0.4. On the other hand, the predicted ΔF/F values of high response sequences (S4-S6) are always underestimates by a difference of 0.4 to 0.8.

**Figure 3.**
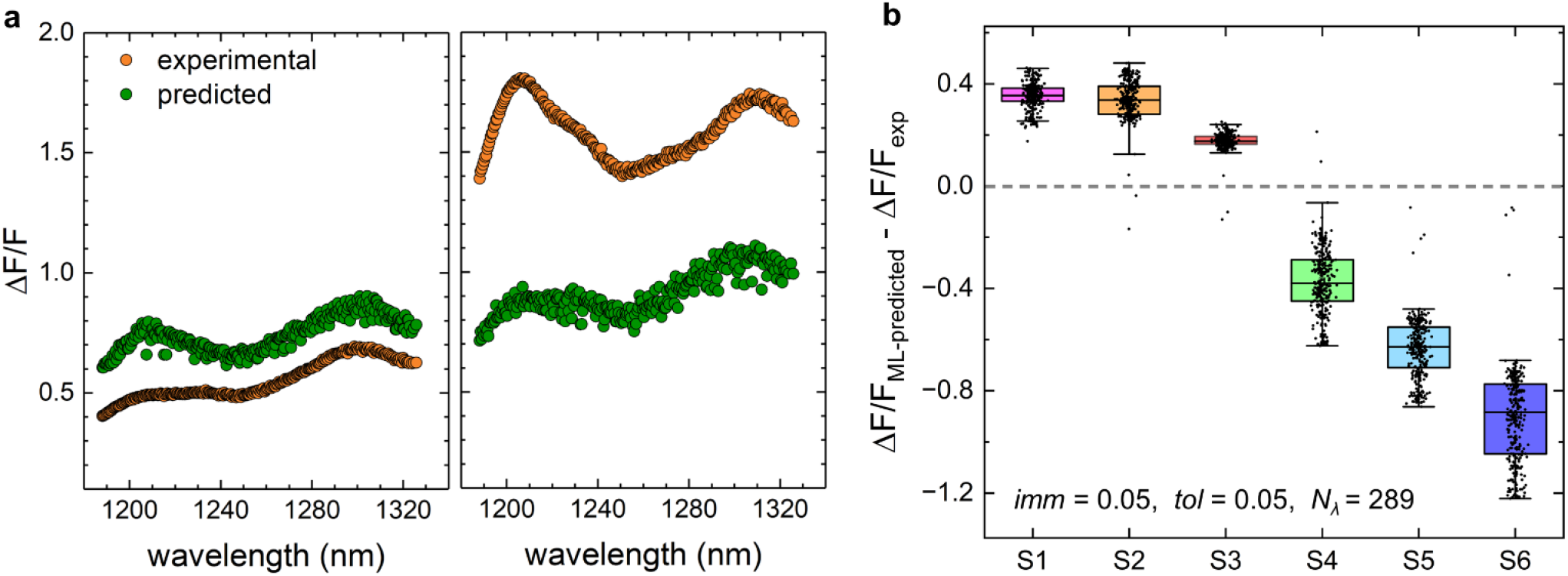
Predicting ΔF/F values for DNA-SWNT conjugates at selected wavelengths using SVM regression models. a) Comparison of measured and predicted ΔF/F values for representative low response (S1, left) and high response (S4, right) sequences. ΔF/F values shown in green are predictions made from a single SVM regression model. b) Differences between the experimentally measured and the predicted ΔF/F values for six testing sequences S1 – S6. The points shown for each sequence represent the differences obtained from five independent SVM regression models (r^2^ > 0.4) at 289 selected wavelengths. The model parameters are defined in the inset.

### 3.3 Procedure for predicting new DNA-SWNTs with high ΔF/F response to serotonin from distributions of predicted ΔF/F values at selected wavelengths

After training the SVM regression models at the selected wavelengths, we used them to predict ΔF/F values for six testing sequences S1 – S6. The violin plots of all the predicted ΔF/F values for the six testing sequences are shown in **Figure** To determine the model parameters *imm* and *tol* that result in the most useful models, we examined the behavior of the distribution plots for *imm* values of 0, 0.05, 0.1, 0.15 and 0.2, and *tol* values of 0.05 and 0.1. Since some choices of parameters lead to fewer than five models with r^2^ > 0.4 for some of the selected wavelengths, the number of wavelengths for which predictions are made varies (from 62 to 289).

The violin plots in **Figure 4** strongly vary in their widths, i.e. the range covered by the predicted ΔF/F values, in dependence of *imm* and *tol* parameter values. Some choices of *imm* and *tol* lead to broad violin plots that span a large range of ΔF/F values, such as when *tol* = 0.05, or when *tol* = 0.1 and *imm* values are large. For such wide violin plots, determining the nature of the response of the tested sequence is difficult, since the distributions in the violin plots overlap, and ΔF/F_ML-predicted_ values forming the distributions span a wide range of values, e.g. between 0.3 and 1.2. Despite overlapping distributions of ΔF/F values of sequences S1 – S6, most of the p-values examining these distributions are < 0.05, indicating that most of the distributions are statistically different (**Figure S3**). With such wide distributions, it becomes difficult to discriminate by visual analysis between sequence responses and to provide with high confidence new sequences which are likely to be either high or low response. However, the visual analysis can be used in conjunction with p-values to identify sequences with outlier distributions of ΔF/F values.

**Figure 4.**
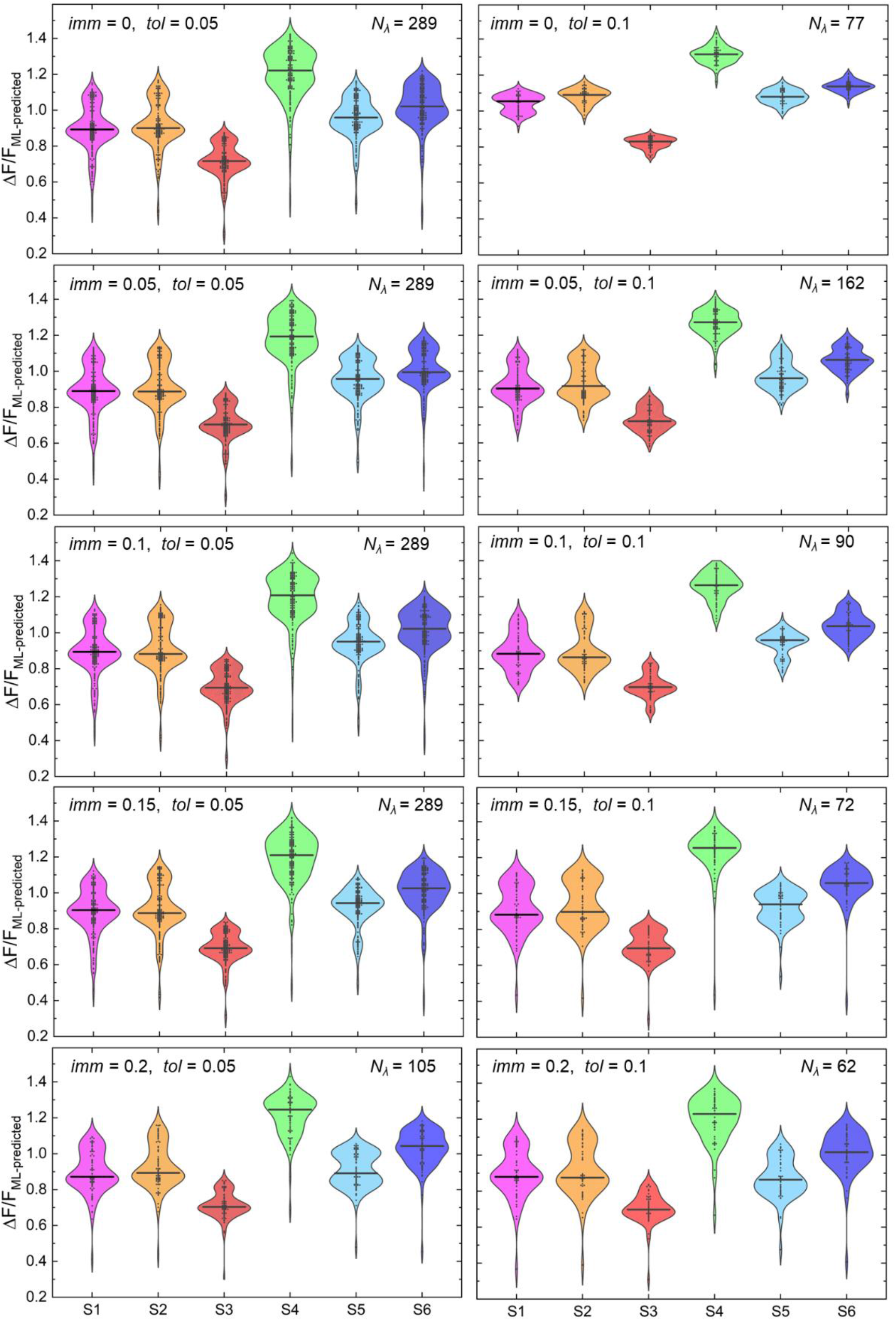
Distributions of predicted ΔF/F values for S1 – S6 sequences, shown as violin plots, where ΔF/F values predict the response of DNA-SWNT conjugates to serotonin. ΔF/F values for each sequence are obtained from multiple wavelengths and from five distinct SVM regression models. The number of wavelengths contributing to the distributions, *N*_*λ*_, and the parameters used in dataset curation, *imm* and *tol* are reported in each plot. The vertical black lines in each violin plots indicate median values of ΔF/F_ML-predicted_.

The violin plot widths are narrowest when *imm* is set to 0 and *tol* is set to 0.1, and it is for such narrow distributions that we can most easily examine for which sequences the ΔF/F value distributions form outliers, as seen if **Figure 4** (top right). For these settings of *imm* and *tol* parameters, the visual analysis allows us to identify S3 as a low response outlier, and S4 as a high response outlier. The p-value analysis (**Figure S3**) also confirms that sequences S3 and S4 are statistically different from the other sequences for these settings of *imm* and *tol* parameters. On the other hand, S1, S2, S5 and S6 sequences are predicted to have similar responses by our models, and their distributions of ΔF/F values overlap. These results show that our models can predict only some new sequences that will have low or high response in experiments, but our models do not have enough knowledge to predict many other possible sequences. This model deficiency is likely due to the relatively small size of the training dataset, compared to the large size of the space of all possible 18-nt long DNA sequences that can be chosen (4^18^ ~ 69 x 10^9^). Therefore, our models are likely to miss many high / low response sequences in that complete sequence space. However, they may be used to predict a subset of the useful sequences that can be then experimentally tested and may lead to the discovery of new DNA-SWNT sensors with the desired high or low response to the selected analyte. Overall, our results confirm that ensembles of ML models trained to predict ΔF/F values of DNA-SWNTs at multiple wavelengths can predict and distinguish some DNA-SWNT conjugates with significantly different high / low response compared to other possible DNA-SWNT conjugates.

After examining the distributions of ΔF/F values predicted at multiple wavelengths for S1 – S6 sequences, **Figure 5** compares one of these distributions to ΔF/F values predicted at a single wavelength (1195 nm), as done in our previous work^17^. Both individual predictions and the distributions of predictions exhibit similar trends, with sequence S3 showing the smallest predicted ΔF/F values, and sequence S4 exhibiting the largest predicted ΔF/F values. While the trends remain consistent, the distributions offer a broader range of predicted ΔF/F values. Utilizing such distributions may enhance confidence in predictions, facilitating the selection of new sequences for experimental examination.

**Figure 5.**
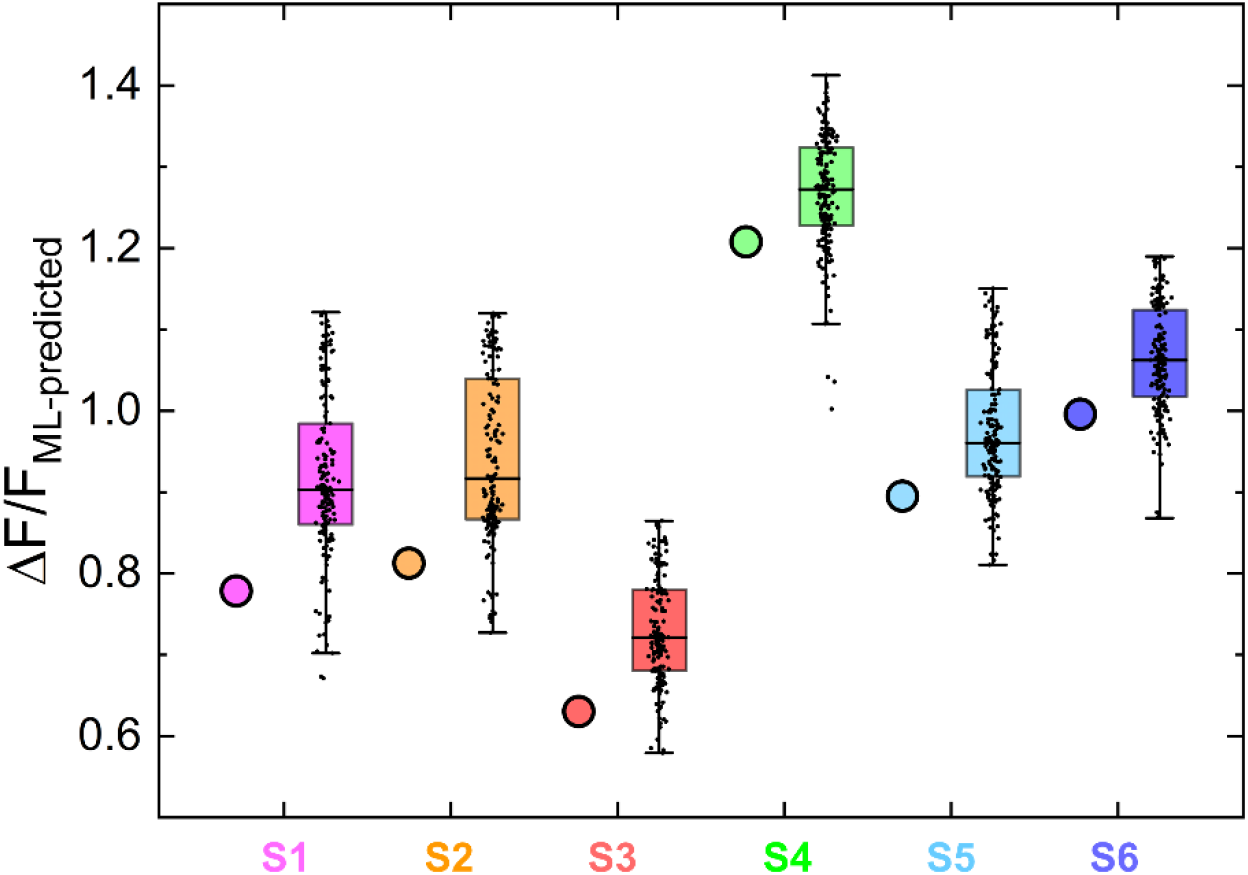
Comparison of ΔF/F values predicted at a single wavelength (1195 nm) and distributions of ΔF/F values predicted at multiple wavelengths for S1 – S6 sequences, shown as boxplots. The reported ΔF/F values predict the response of DNA-SWNT conjugates to serotonin. The values are shown for *imm* = 0.05 and *tol* = 0.1, where the number of wavelengths contributing to the distributions, *N*_*λ*_, is 162.

## 4. Conclusion

In this work, we built upon our prior success in machine learning approaches, which led to the prediction of five novel DNA-SWNT sensors with superior responses to serotonin compared to any DNA-SWNT in the original dataset. Our previous ML models, while effective, were trained solely on responses of DNA-SWNTs to serotonin at a single wavelength (1195 nm) extracted from complete fluorescence emission spectra of the tested sample. Notably, the information residing in the remaining parts of the spectra was left untapped for ML predictions. Here, we address this limitation by leveraging information from multiple wavelengths across all spectra obtained experimentally. Our analysis of the whole spectra of all the DNA-SWNT conjugates in the dataset lead to the first important insight. The crucial role of DNA sequence suggests the potential existence of distinct binding modes of DNA molecules to the target analyte (here, serotonin) in the presence of a hydrophobic SWNT surface. Notably, our observations suggest that the SWNT chirality plays a less substantial role in influencing analyte binding.

Our ML models trained in the present work predict ΔF/F response at over a hundred wavelengths for each sequence in the dataset of the tested DNA-SWNT conjugates. These predictions are then statistically analyzed to create a distribution of ΔF/F predictions for a given sequence. We evaluate the performance of this novel approach, which utilizes data from broader regions of experimental spectra, providing a comprehensive examination compared to the methodology outlined in Ref^17^. While our new machine learning (ML) models exhibit success in predicting specific high-response DNA sequences, some limitations are also apparent. The results demonstrate that our models, despite their efficacy, are constrained by their inability to comprehensively predict all potential sequences with either low or high response. This limitation stems from the relatively modest size of the training dataset in comparison to the vast space of all possible 18-nt long DNA sequences (~ 69 x 10^9^). Consequently, our models might overlook numerous high- or low-response sequences in the complete sequence space. Nevertheless, they offer a valuable tool for predicting subsets of sequences that can be experimentally tested, potentially leading to the discovery of novel DNA-SWNT sensors with desired response profiles to specific analytes.

It may be advantageous for future studies to incorporate more experimental data encompassing whole spectra as inputs for training ML models, as compared to using only single datapoints from each spectrum. This approach holds promise for generating distributions of predicted spectral response values, thereby increasing confidence in predictions, and guiding the selection of new systems for experimental testing. This refinement in model input has the potential to enhance the robustness of our predictions and facilitate the identification of sequences with specific response characteristics. Notably, the experimental workflow upon which our analysis is based, and the ML models developed here, are analyte-agnostic. Therefore, this work could seed rapid discovery of DNA-SWNT nanosensors for a large range of analytes. In summary, our new approach may be useful for future attempts to predict spectra of different chemical systems and to discover new DNA-based systems with a desired optical response to introduced perturbations.

## ASSOCIATED CONTENT

### Supporting Information

A table of DNA sequences in DNA-SWNT conjugates within our dataset, and the assigned sequence identification numbers; analysis of minimum and maximum ΔF/F values and the difference between the two values in our dataset, a schematic definition of parameters used to process and curate our datasets, distributions of ΔF/F values at several wavelengths across all DNA-SWNT conjugates in our dataset, a table of six DNA sequences (S1 – S6) selected for independent validation of machine learning models, a comparative p-value analysis of predicted ΔF/F values for S1 – S6 sequences using heatmaps.

### Data and Software Availability

All methods described in this section were implemented using the Python programming language. The associated code is available for public access and can be found at the following GitHub repository: github.com/vukoviclab/PySpectrotonin. The datasets of ΔF/F values at all the measured wavelengths for 136 DNA-SWNT sequences is available at: https://github.com/vukoviclab/PySpectrotonin/blob/main/full.csv.

### Funding

We acknowledge the support of the NSF CBET-2106587 award (to M.P.L. and L.V.) and the computer time provided by the Texas Advanced Computing Center (TACC).

---

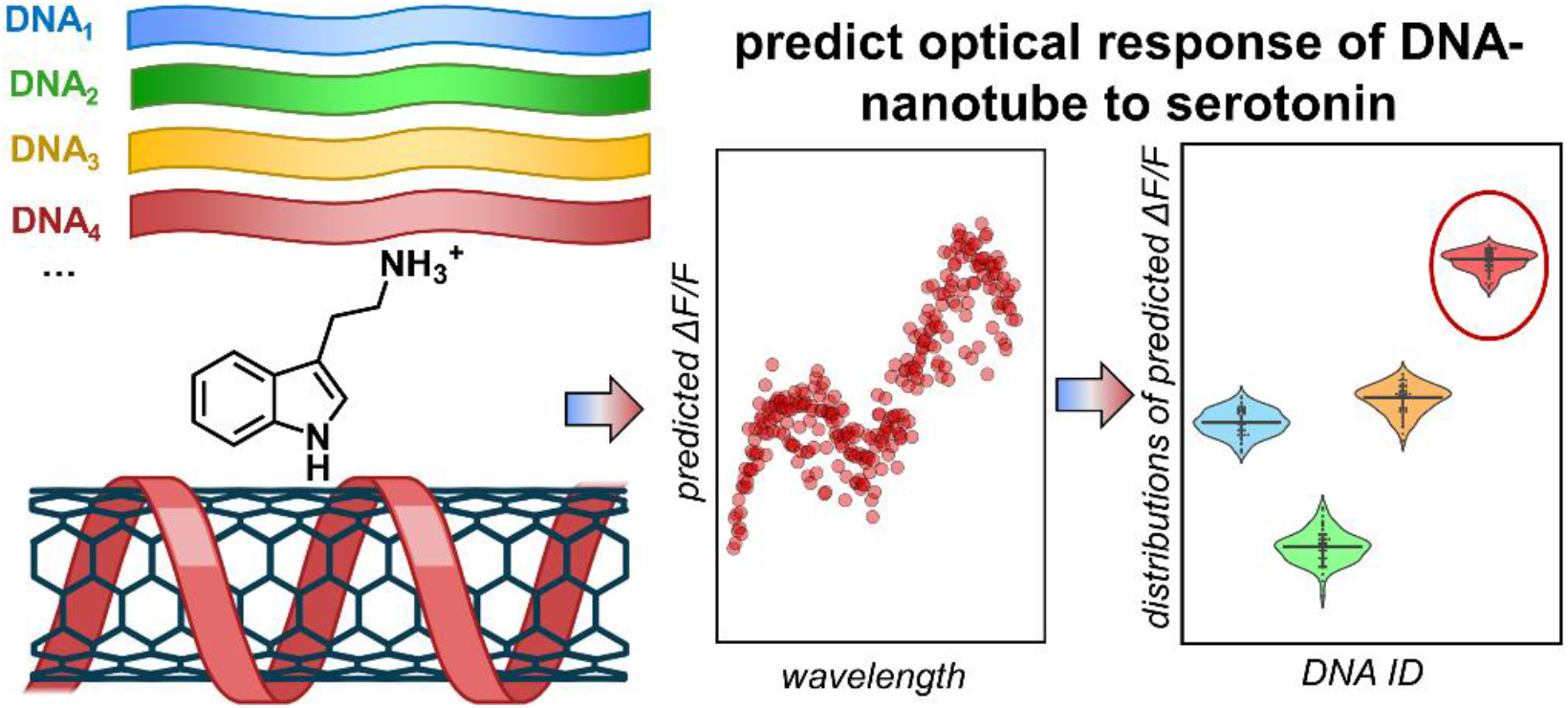

TOC graphic.

