## Supplementary Information for "Predicting Serotonin Detection with DNA-Carbon Nanotube Sensors Across Multiple Spectral Wavelengths"

**Table S1.** DNA sequences in DNA-SWNT conjugates within our dataset, and the assigned sequence identifications (IDs). Six sequences selected for independent evaluation of our models are highlighted. The sequences corresponding to IDs 39 and 115 are duplicates and were removed from the machine learning training dataset to ensure data integrity and the reliability of the training process.

| ID | DNA sequence | ID | DNA sequence |
|---|---|---|---|
| 1 | ACCTGACACGATCCTATG | 44 | TAGCACAGGTCGTCTATT |
| 2 | GGCACAACGCTCGATGCT | 45 | GCCAATATAGCCCTTCCG |
| 3 | ATTACAGCGGACAAGTGT | 46 | AATCACTGCAATGGTCGT |
| 4 | TAAGGCCGATCCCACTAT | 47 | AACACATTGACGTGCACT |
| 5 | TGACTCCATAACAGTGTG | 48 | GGGCTGTGCCGTCATGCG |
| 6 | GACACCCTGGACCCGTCG | 49 | GATGGGGAATCATGCGTG |
| 7 | TGGCGTACAAACCGTCTG | 50 | ACACAGCATCATTCCGCT |
| 8 | ACACACTCTACTCTTCCA | 51 | GCACCAACCAGCCGTCTG |
| 9 | GACGTTGTGCCCAAGTTG | 52 | TCACCACATTCGACGGCG |
| 10 | AAGGGACTGAAAGCAATG | 53 | ACCACAAGTGACTGTCCT |
| 11 | GATCCAACCGCTGCCACA | 54 | GCCGACATGACTCCTCCT |
| 12 | ACGACGTACACTCCTCCT | 55 | ACACACCAATGACCTGTG |
| 13 | AACCGCATGTACTCTCCG | 56 | TACCCACACCACACACTG |
| 14 | AACATGCACAGACGTCCG | 57 | ACTGCACATCGACGCGCG |
| 15 | AACCATGCACAACGCGTG | 58 | ATTGCCGCCATCCTCATG |
| 16 | ACACAACCTGCTCCTCCT | 59 | AGGCCACCGTCGCACGTG |
| 17 | CCCCCCCCCCCCCCCCCC | 60 | AACACCACACACGGCGCT |
| 18 | ACGCACAATCCGGCACTT | 61 | AGCACACTCCACTCCGCT |
| 19 | ACAGACTGCAGTCATGTG | 62 | GCACACACCAGCCGTCTG |
| 20 | ACACCAGCCACACGTGCG | 63 | AACCACACACCGTCCGCT |
| 21 | ACGCACCGACAGCACACT | 64 | ACCACACCATCGACGCGT |
| 22 | ACACCACACCACACCGAT | 65 | AGCCACACGACGCGCTCT |
| 23 | ACGACAACCAACACTGTG | 66 | ACGGCACACACCATCGCT |
| 24 | AGCACACTACACACGGCG | 67 | ACGACACTGCACGACGCG |
| 25 | ACACCACCTCACGACGTG | 68 | ACGGCAACTCCCATTCCG |
| 26 | ACACCACCAGACACTGCG | 69 | ACGACACCACACTGCTCT |
| 27 | ACCAACACCAGCCGTGCG | 70 | ACCGCATCGACATGTGCT |
| 28 | ACACACACCACACGTGCT | 71 | ACCGCACGAGCCAGTGTG |
| 29 | ACACAACACCCGACGCGG | 72 | TCACCACATTCCGCTGTG |
| 30 | ACACACACAACGACGCGG | 73 | ACCGAGAGCAGACGATGT |
| 31 | AGCACAACACGGCAACCT | 74 | GCAGCGTGACTTGACGTG |
| 32 | AACACACCACAGACTCTG | 75 | AACACGGCCCTCATGTCG |
| 33 | ACACACCATCAGACGCCG | 76 | AGCCGTATGCACACCTCA |
| 34 | AGCAGCACACGACACACT | 77 | ACACACCGTTCATCCGCG |
| 35 | ACGCCAACACATTCCGCT | 78 | GCTGATCGACGACACGTG |
| 36 | AACACACACAGCCGTCCG | 79 | ACACCACAGCACTCCGAT |
| 37 | AACACACACAGACGCACG | 80 | ACACCCAACGTCTGCTCT |
| 38 | AGCACCAGACAGCACACT | 81 | ACACCCTAACTCCGCTCT |
| 39 | ACCACGATCCTCACTCCG | 82 | ACACCCTGAGTCCGCACA |
| 40 | ACAGACCGACGTGTGCTG | 83 | ACACACCGATCCACCGCT |
| 41 | TGGGAGCCATCTTGTGCG | 84 | ACATACCCACTCCGCTCG |
| 42 | GTTCAGCCTTTTCGTTCG | 85 | GCACACCGATCCTACCAG |
| 43 | GGAATCTCCGGCGTCTAT | 86 | ACACACCCTAATCTCGCT |

| ID | DNA sequence | ID | DNA sequence |
|---|---|---|---|
| 87 | ACAAACCGCTCATCCGAT | 124 | GACCACTCCAATTCCGCT |
| 88 | ACACTCCGACCCTTCTCG | 125 | AGCACCACCAGACTCCTG |
| 89 | AACGCCACCCTAACTCCG | 126 | ACGCCACACCATTCCGCT |
| 90 | AGCCCGAACCAGACACCG | 127 | AACCCGAAGCCTGGACCT |
| 91 | AACCCGAACCTAACTGCG | 128 | AGCACAACACGGCACCGT |
| 92 | AACGCAACACGACCTGTG | 129 | ACAACAACACCTTCCGCT |
| 93 | AACCCAGACCGACCACCT | 130 | AACACAACAGCTCCTCCT |
| 94 | GACCCAAAGCCAACACCT | 131 | AAGGCAACCAGACGTCCG |
| 95 | GACCCTAACACAGCACCA | 132 | ACAACCACCGATCCATCG |
| 96 | AACCCTAACCGATCACTG | 133 | ACACTCTCCCATTCCGCT |
| 97 | AACACGACCCGACCTGTG | 134 | AACACCACTCGACGCGTA |
| 98 | GACCCAAACCTACCTCCA | 135 | AGCCAACATCATTCCGCT |
| 99 | TACCACTCCAACTCCGCT | 136 | ACACCCACTCCACACGCT |
| 100 | ACACGCCCTAACTCCGCT | 137 | GACCCACACCAACCAGTG |
| 101 | AGGCACACTCATCCGCGT | 138 | AGCACCCTCTACAGCACA |
| 102 | ACACCGCCTAATGCACCT | | |
| 103 | CACGATCCTACTCACGCT | | |
| 104 | ACACCCGATCCTGCACCG | | |
| 105 | AACCAACATCCTTCCGCT | | |
| 106 | GGCAACGATACTCAACCT | | |
| 107 | ACCACACCTCATTCCGCT | | |
| 108 | AACACCGATCCTCCATGT | | |
| 109 | ACACACAGACTCACCGCT | | |
| 110 | ACACCGATCCTACGCACT | | |
| 111 | GCCACCAGACTCAATGCT | | |
| 112 | ACACCGATCCTAACTCCG | | |
| 113 | AACACAACATTCCGCTCG | | |
| 114 | AACCTCACACTCATTCCG | | |
| 115 | ACCACGATCCTCACTCCG | | |
| 116 | ACGACCCTATGAACCTCG | | |
| 117 | GACCACAGAATACGGCCT | | |
| 118 | AGCAACAGACGACCTGCT | | |
| 119 | AGCACAGCACGACGCGTA | | |
| 120 | AACACCGCACCATCCGAT | | |
| 121 | AACGACACACATTCCGCT | | |
| 122 | AACCCGACACCACACCTG | | |
| 123 | ACCACATCCACAGCCGAT | | |

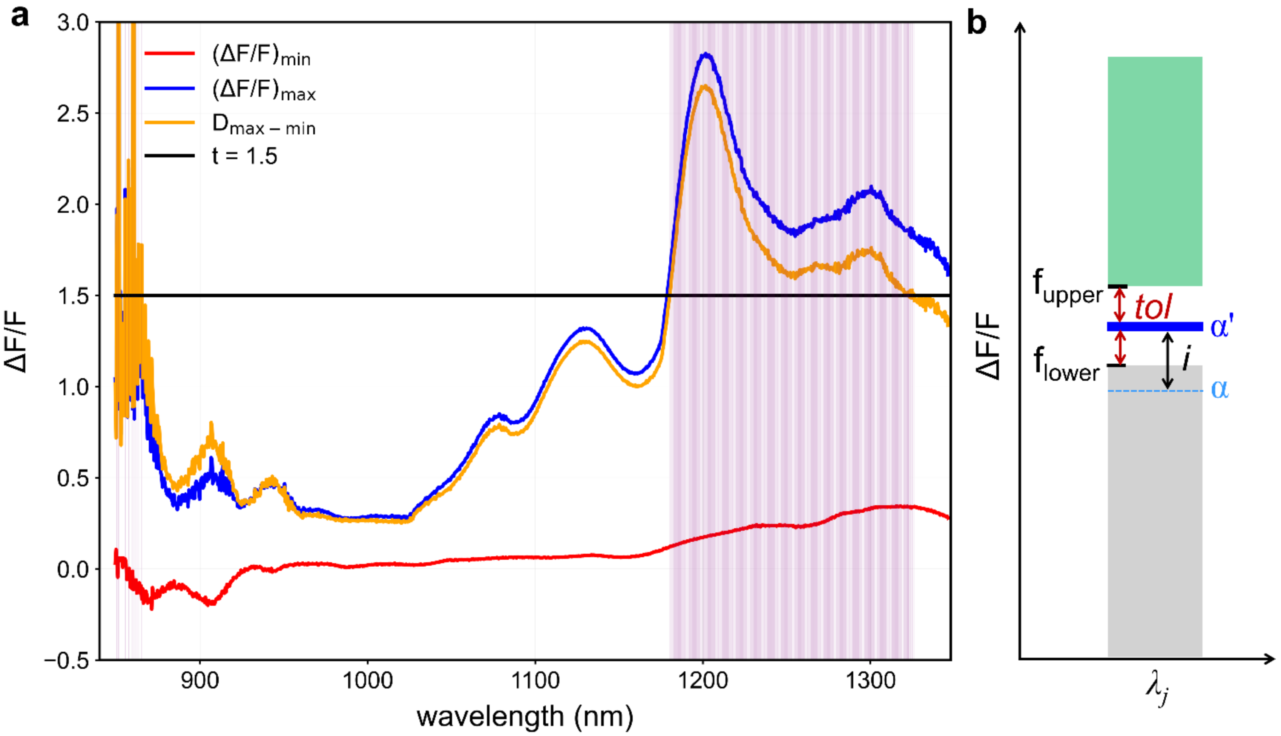


**Figure S1**. a) The analysis of the range of ΔF/F values for 130 DNA-SWNT conjugates in the experimentally-obtained dataset. The plot shows the minimum ΔF/F values across the conjugates (red), the maximum ΔF/F values across the conjugates (blue) and the difference between two values, $D_{max\text{-}min}$, at each wavelength (orange). The wavelengths for which $D_{max\text{-}min} > t = 1.5$, are highlighted with purple shading. b) A diagram demonstrating the range of ΔF/F values used to remove datapoints from the original dataset and achieve bimodal-like distribution, as described in Methods. All the parameters shown in the panel are defined and described in Methods.

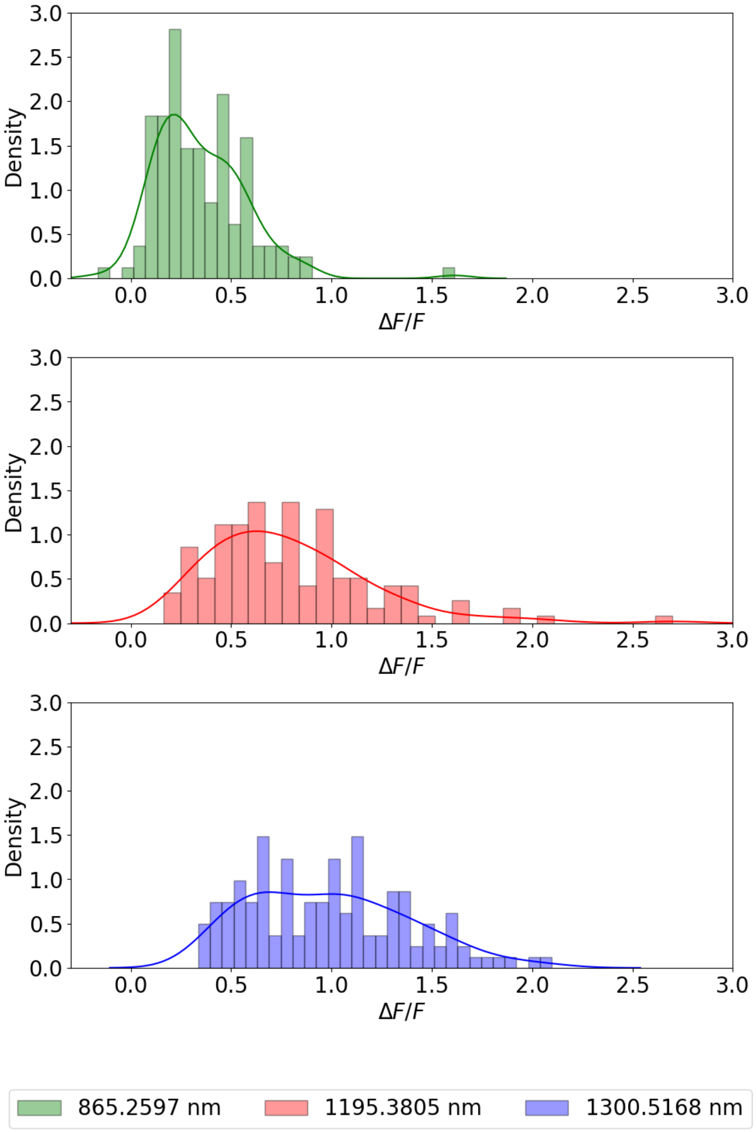


**Figure S2.** Distributions of ΔF/F values for 136 DNA-SWNT conjugates at three selected wavelengths, 865.3 nm (green), 1195.4 nm (red), and 1300.5 nm (blue).

**Table S2.** Six DNA sequences selected for independent validation of the ML models.

| ID | DNA sequence | index | color |
|---|---|---|---|
| 47 | AACACATTGACGTGCACT | S1 | |
| 59 | AGGCCACCGTCGCACGTG | S2 | |
| 42 | GTTCAGCCTTTTCGTTCG | S3 | |
| 31 | AGCACAACACGGCAACCT | S4 | |
| 35 | ACGCCAACACATTCCGCT | S5 | |
| 38 | AGCACCAGACAGCACACT | S6 | |

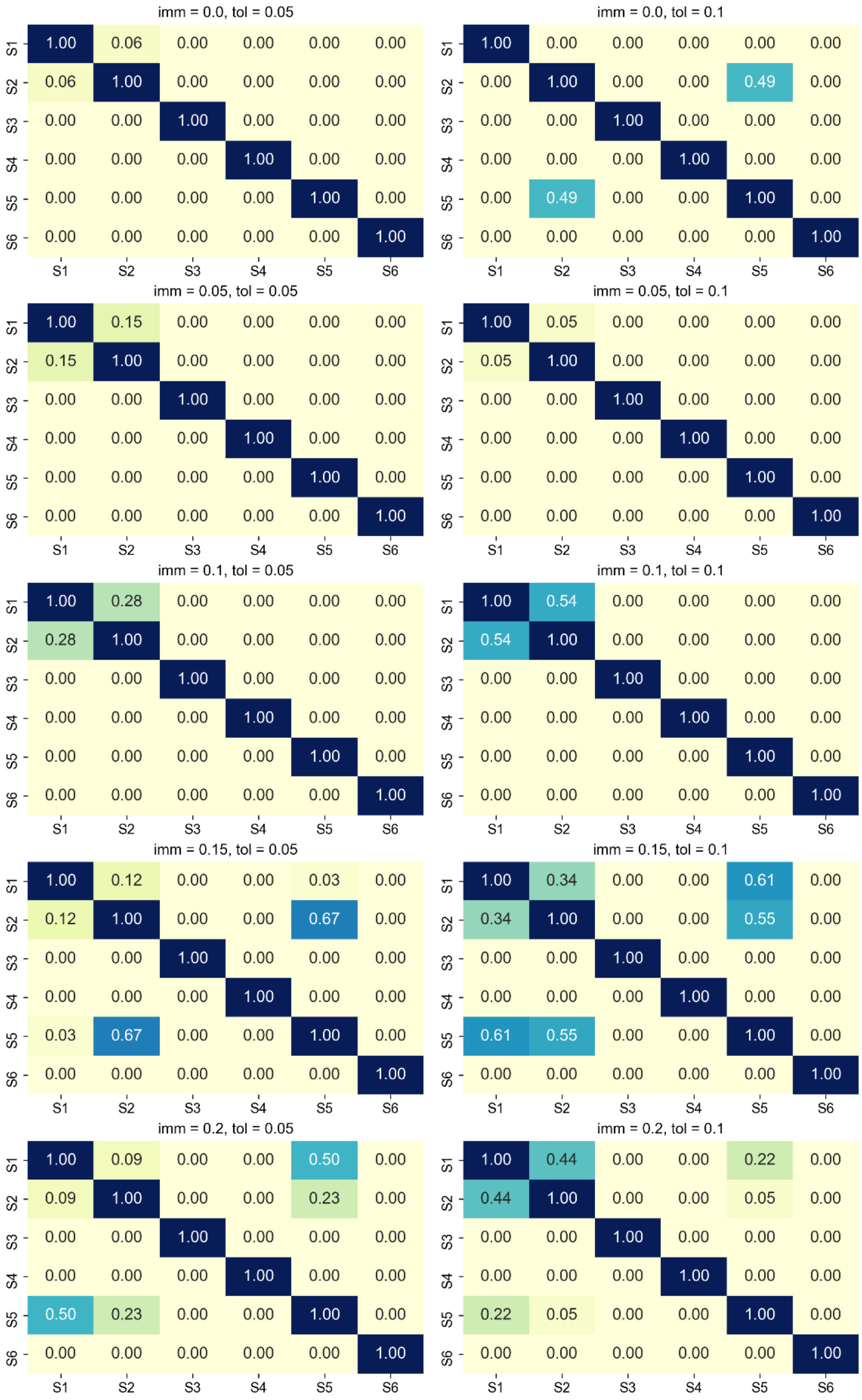


**Figure S3.** Comparative p-value analysis of predicted ΔF/F values at all retained wavelengths for S1 – S6 sequences, for ML regression models trained using different values of *imm* and *tol* parameters. Each cell within the heatmaps contains a p-value resulting from a two-tailed t-test, reflecting the statistical comparison between each pair of conditions.
